# Reverse-engineering Recurrent Neural Network solutions to a hierarchical inference task for mice

**DOI:** 10.1101/2020.06.09.142745

**Authors:** Rylan Schaeffer, Mikail Khona, Leenoy Meshulam, International Brain Laboratory, Ila Rani Fiete

## Abstract

We study how recurrent neural networks (RNNs) solve a hierarchical inference task involving two latent variables and disparate timescales separated by 1-2 orders of magnitude. The task is of interest to the International Brain Laboratory, a global collaboration of experimental and theoretical neuroscientists studying how the mammalian brain generates behavior. We make four discoveries. First, RNNs learn behavior that is quantitatively similar to ideal Bayesian baselines. Second, RNNs perform inference by learning a two-dimensional subspace defining beliefs about the latent variables. Third, the geometry of RNN dynamics reflects an induced coupling between the two separate inference processes necessary to solve the task. Fourth, we perform model compression through a novel form of knowledge distillation on hidden representations – Representations and Dynamics Distillation (RADD)– to reduce the RNN dynamics to a low-dimensional, highly interpretable model. This technique promises a useful tool for interpretability of high dimensional nonlinear dynamical systems. Altogether, this work yields predictions to guide exploration and analysis of mouse neural data and circuity.

## 1 Introduction

Decision making involves weighing mutually-exclusive options and choosing the best among them. Selecting the optimal action requires integrating data over time and combining it with prior information in a Bayesian sense. Here we seek to understand how RNNs perform hierarchical inference. For concreteness and for the later goal of comparing against the mammalian brain, we consider a perceptual decision-making and changepoint detection task used by the International Brain Laboratory (IBL) [1], a collaboration of twenty-two experimental and theoretical neuroscience laboratories. Optimally solving the IBL task requires using sensory data to infer two latent variables, one cued and one uncued, over two timescales separated by 1-2 orders of magnitude.

We address two questions. First, how do RNNs compare against normative Bayesian baselines on this task, and second, what are the representations, dynamics and mechanisms RNNs employ to perform inference in this task? These questions are of interest to both the neuroscience and the machine learning communities. To neuroscience, RNNs are neurally-plausible mechanistic models that can serve as a good comparison with animal behavior and neural data, as well as a source of scientific hypotheses [15, 16, 31, 5, 27, 10, 8]. To machine learning, we build on prior work reverse engineering how RNNs solve tasks [30, 28, 16, 15, 3, 20, 14, 19], by studying a complicated task that nevertheless has exact Bayesian baselines for comparison, and by contributing task-agnostic analysis techniques.

The IBL task is described in prior work [29], so we include only a brief summary here. On each *trial*, the mouse is shown a (low or medium contrast) stimulus in its left or right visual fields and must indicate on which side it perceived the stimulus. Upon choosing the correct side, it receives a small reward. Over a number of consecutive trials (a *block*), the stimulus has a higher probability of appearing on one side (left stimulus probability *p_s_*, right stimulus probability 1 − *p_s_*). In the next block, the stimulus side probabilities switch. The change-points between blocks are not signaled to the mouse. This task involves multiple computations, elements of which have been studied under various names including change-point detection [2, 22].

## 2 Methods

### 2.1 IBL task implementation

Each session consists of a variable number trials, indexed *n*. Each trial is part of a block, with blocks defining the prior probability that a stimulus presented on the trail is shown on the left versus the right. The block side on trial n, denoted *b_n_* ∈ { − 1,1} (−1: left, 1: right), is determined by a 2-state semi-Markov chain with a symmetric transition matrix. The probability of remaining on the same block side as in the last trial is 1 − *p_b_*; the probability of switching block sides is p_b_. The process is semi-Markov because *p_b_* varies as a function of the current block length (*l_n_*) to ensures a minimum block length of 20 and maximum block length of 100, with otherwise geometrically distributed block lengths.

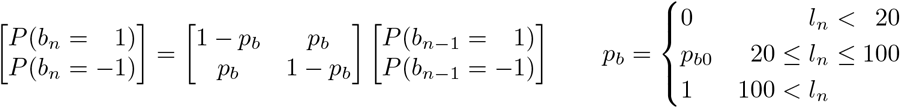

The stimulus (*s_n_* ∈ {−1,1}) presented on trial n is either a left or right stimulus, determined by a Bernoulli process with a single fixed parameter *p_s_*, which gives the probability that the stimulus is on the same side as the current block (termed a *concordant* trial). The probability of a *discordant* trial (stimulus on opposite side of block) is 1 − *p_s_*. In the IBL task, *p_b0_* = 0.02 and *p_s_* = 0.8.

Neural time-constants (10-100 ms) are much shorter than the timescale of trials (~ 1 s), so we model a trial as itself consisting of multiple timesteps indexed by *t*. A trial terminates one timestep after the RNN takes an action (explained in the next subsection) or after timing out at *T_max_* steps, whichever comes first. At the start of the trial, the stimulus side *s_n_* and a stimulus contrast strength *μ_n_* are sampled (Fig. 1). Within a trial, on each step, the RNN receives three scalar inputs. On the first step, all three are 0. For each subsequent step, the RNN receives two noisy observations 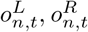, sampled i.i.d. from two univariate Gaussians with mean *μ_n_* for the stimulus side and mean 0 for the other. The third input is a reinforcement signal *r_n,t_*, which takes one of three possible values: a small waiting penalty (−0.05) in every timestep, a reward (+1) if the correct action was taken on the previous step, or a punishment (−1) if the incorrect action was taken on the previous step or the model timed out.

**Figure 1:**
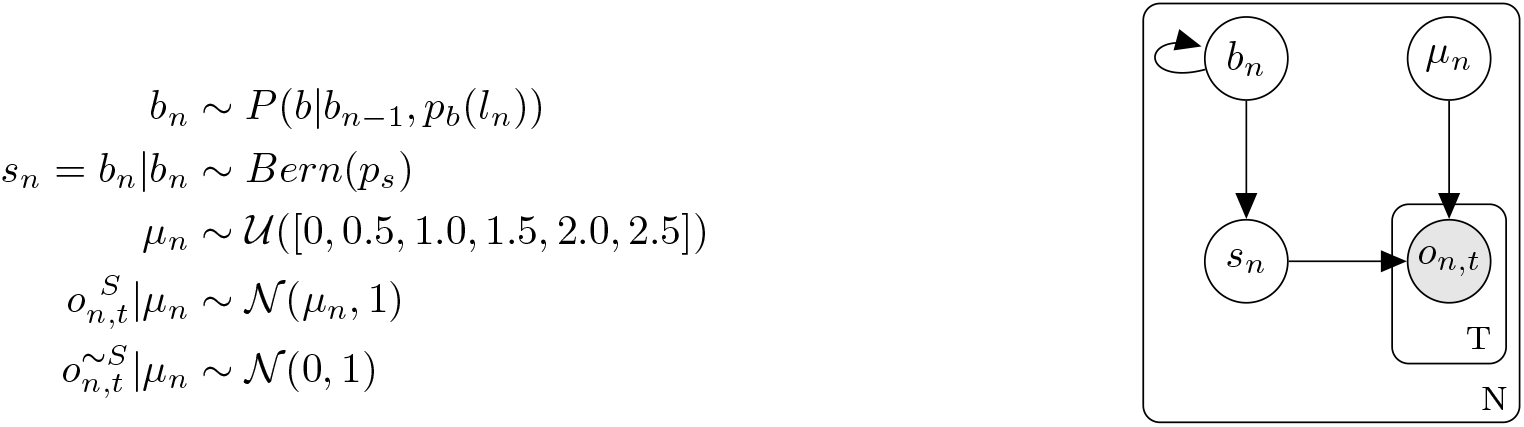
Generative model of the IBL task. Block side *b_n_* is determined by a 2-state semi-Markov chain. Stimulus side *s_n_* is either *b_n_* with probability *p_s_* or *−b_n_* with probability 1 − *p_s_*. Trial stimulus contrast *μ_n_* determines the observations on each timestep *t* within a trial: 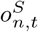 for the stimulus side, 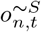 for the non-stimulus side.

### 2.2 Recurrent network architecture and training

On each step, the observation 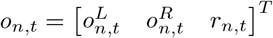 is input to the RNN. Letting *h_n,t_* denote the RNN state on the nth trial and the tth step within the trial, the state is defined by the typical dynamics

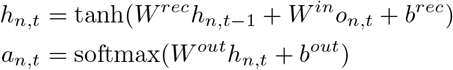

where *a_n,t_* is a probability distribution over the two possible actions (left or right). An action is defined by when the probability mass on either action exceeds a fixed threshold (0.9). We present RNNs with 50 hidden units, but the results are similar for other numbers of units (e.g. 100, 250). We train the RNN under cross entropy loss using stochastic gradient descent with initial learning rate 0.001 and initial momentum = 0.1. RNN parameters were initialized using PyTorch defaults. PyTorch and NumPy random seeds were both set to 1. Our code will be publicly available at https://github.com/int-brain-lab/ann-rnns.

### 2.3 Normative Bayesian baselines

The IBL task involves inference of two latent variables, the stimulus side and the block side. Exact inference can be decomposed into two inference subproblems that occur over different timescales, which we term *stimulus side inference* and *block side inference*:

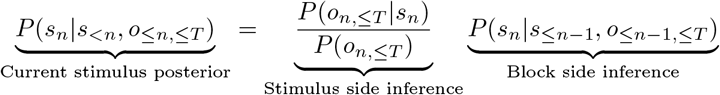

where ·≤*m* denotes all indices from 1 to *m*, inclusive. We consider two Bayesian baselines. The *Bayesian actor* performs the task independently from the RNN, but using the same action rule (i.e. an action taken when its stimulus posterior passes the action threshold). The *Bayesian observer* receives the same observations as the RNN, but cannot decide when to act; the RNN therefore determines how long a trial lasts. The Bayesian actor tells us what ceiling performance is, while the Bayesian observer tells us how well the RNN could do given when the RNN chooses to act.

Other than this difference, the Bayesian actor and the Bayesian observer are identical. Both assume perfect knowledge of the task structure and task parameters, and both are comprised of two separate submodels performing inference. The first submodel performs stimulus side inference given the block side, while the other submodel infers block changepoints given the history of true stimuli sides. True stimuli sides can be determined after receiving feedback because the selected action and the ensuing feedback signal (correct or wrong) together fully specify the true stimulus side.

*Stimulus side inference* occurs at the timescale of a single trial. Since observations within a trial are sampled i.i.d., the observations are conditionally independent given the trial stimulus strength μ_n_. The likelihood is therefore:

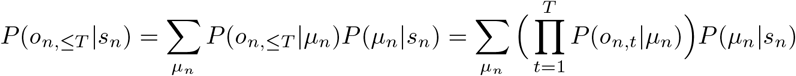

*Block side inference* occurs at the timescale of blocks, based on knowledge of the history of true stimuli sides. Our Bayesian baselines assume that the block transitions are Markov (instead of semi-Markov). Both baselines perform Bayesian filtering [25] to compute the block side posterior by alternating between a joint and a conditional, and normalizing after each trial:

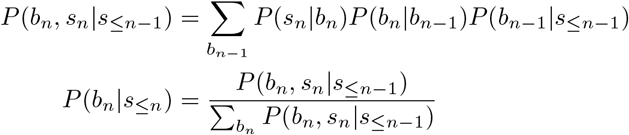

## 3 Results

### 3.1 RNN Behavior

#### 3.1.1 RNN behavior matches ideal Bayesian observer behavior

We start by quantifying the performance of the RNN. Strikingly, the RNN achieves performance nearly matching the Bayesian observer (Fig. 2a); all three agents display similar accuracy as a function of trial stimulus strength *μ_n_*: the fraction of correct actions is highest for strong stimulus contrast and lowest for weak stimulus contrast. Furthermore, the performance of all three agents is well above chance for zero-contrast trials, meaning all three exploit the block structure of the task.

**Figure 2:**
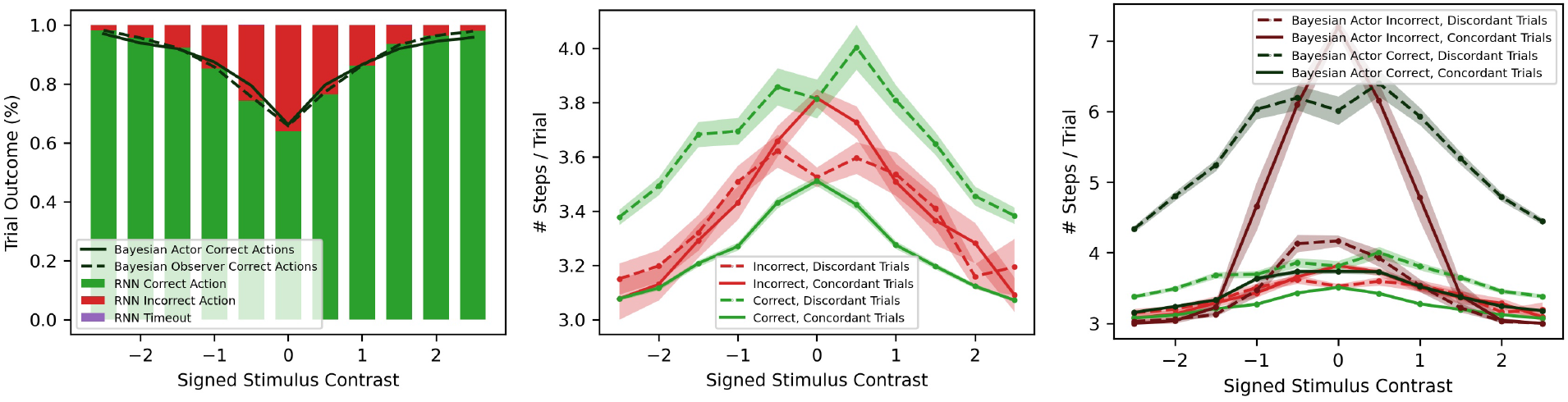
(a) RNN fraction of correct action almost matches Bayesian observer and Bayesian actor. (b) RNN chronometric curves show longer integration time on low contrast trials. (c) Bayesian actor chronometric curves show the actor responds significantly more slowly on low contrast trials than the RNN. RNN curves from (b) have been added for comparison.

Chronometric curves, which quantify how quickly the agents select an action as a function of trial stimulus strength, show that both the RNN and the Bayesian actor respond faster on concordant trials (when the stimulus side matches the block side, a higher probability event) than discordant trials (Figs. 2ab). While both the RNN and the Bayesian agents act more slowly for low trial stimulus contrast, the RNN acts significantly faster than the Bayesian actor on trials with low trial stimulus strength (Figs. 2ab. This suboptimal integration of within-trial evidence by the RNN partly explains its slightly worse performance than the Bayesian actor.

#### 3.1.2 RNN leverages block prior when selecting actions

We next explored to what extent the RNN leverages the block prior to select actions. The RNN and the Bayesian actor both perform much better on concordant trials than discordant trials (Fig. 3a), but the discrepancy shrinks for large stimulus contrast values. Additionally, in both agents, the gap in stimulus side inference for concordant versus discordant trials is greater for low-contrast trials than for high-contrast trials (Fig. 3a), meaning that when the contrast strength is low, the prior dominates the likelihood, while at high contrasts, the likelihood dominates the prior. The RNN’s behavior approaches that of the Bayesian actor, suggesting that the RNN weighs the stimulus side likelihood with the block prior near-optimally.

**Figure 3:**
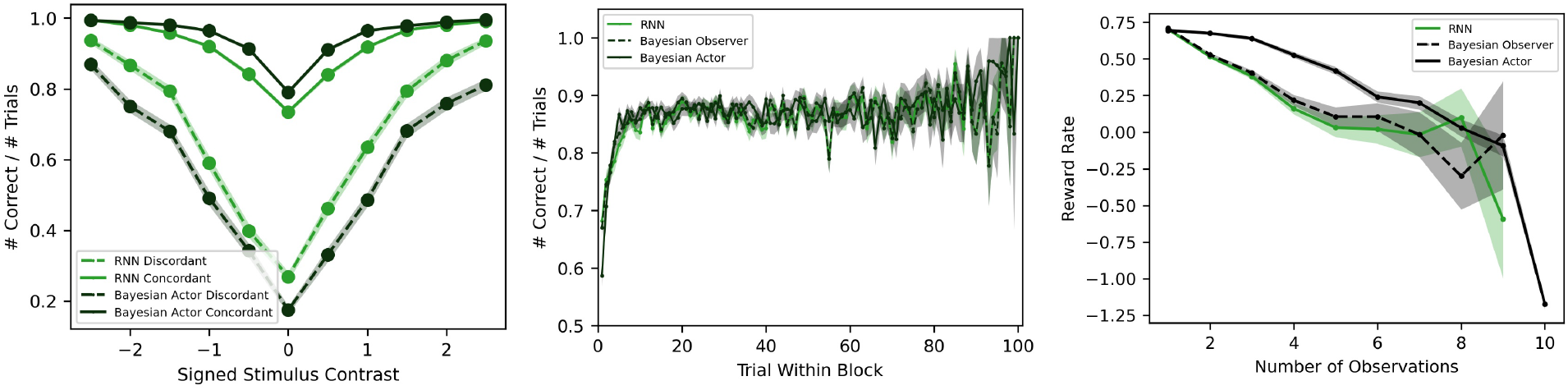
(a) Psychometric curves for RNN and Bayesian actor closely match. Low stimulus contrast values and discordant trial curves indicate the RNN disproportionally weights the likelihood. (b) RNN fraction of correct trials rapidly increases following a block change-point, closely matching the Bayesian observer. (c) The reward rate of the RNN nearly matches the reward rate of the Bayesian observer, but falls short of the Bayesian actor.

There are, however, two small quantitative difference indicating the RNN underweighs the prior. First, the concordant-discordant gap is smaller in the RNN than the Bayesian actor. Second, for zero-contrast stimuli, the Bayesian actor’s accuracy directly reflects the block prior (0.2/0.8), while the RNN’s accuracy is slightly contracted towards chance performance (.5/.5). This is likely not due to deficiencies with inferring the block prior, as the RNN’s fraction of correct answers rapidly climbs following a block change-point (Fig. 3b), closely matching the Bayesian observer and the Bayesian actor, and therefore indicating that change-point detection in the RNN is near-optimal.

### 3.2 RNN Representations

#### 3.2.1 RNN learns 2D dynamics to encode stimulus side and block side

We next sought to characterize how the RNN’s dynamics subserve inference. The first two principal components (PCs) of RNN activity explain 88.74% of the variance, suggesting it has learnt a low-dimensional solution. The RNN readout matrix *W^out^* converts the hidden states *h_n,t_* into actions *a_n,t_*, explicitly giving us the direction along which the RNN encodes its stimulus side belief; we term this the *stimulus readout* vector. 93.39% of the stimulus readout’s length lies in the 2D PCA plane.

To identify how block side is encoded in RNN activity, we trained a logistic classifier to predict block side. This classifier had 82.6% accuracy on 33% heldout test data. A separate classifier for block side trained from only the 2-dimensional PCA plane of RNN activity reached 82.5% accuracy (Fig. 4a). In short, the RNN’s PCA plane encompasses the two latent variables being inferred: these two dimensions are sufficient to decipher how the network solves the task. Importantly, the (right-side) block and (right-side) stimulus readouts are non-orthogonal (subtending an angle of 73° to each other in the high-dimensional RNN space, and 68° in the PCA plane). This deviation from orthogonality is modest but critical to how the network performs hierarchical inference (as we will explain below).

**Figure 4:**
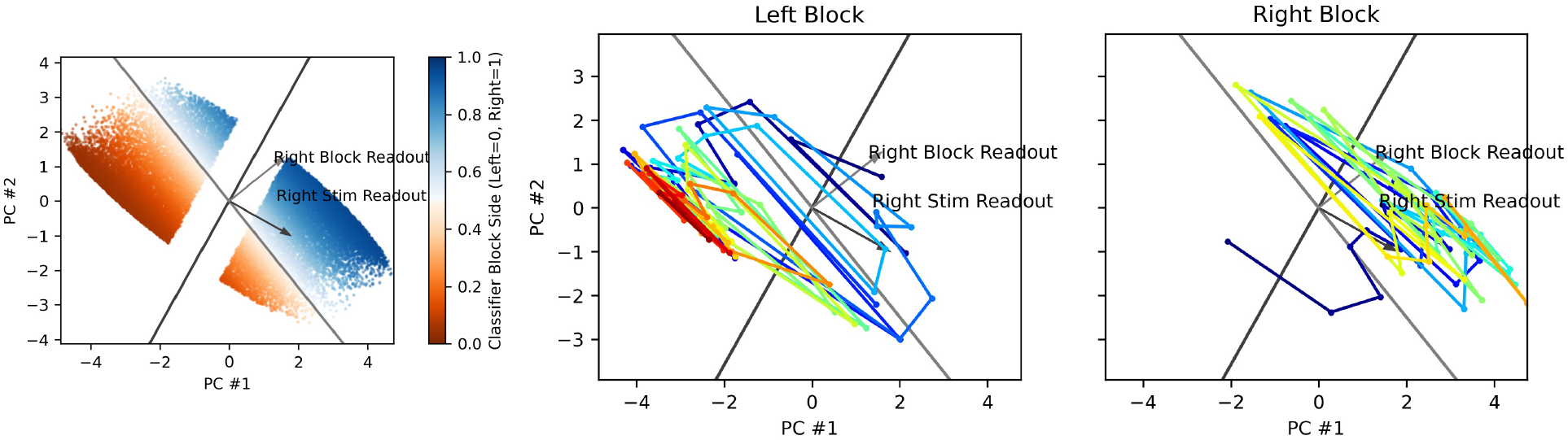
(a) Logistic regression classifies block side from RNN activity with 82.6% accuracy on 33% test data. The block readout and stimulus readout directions are non-orthogonal. (b-c) Example RNN state-space trajectories in a left block and a right block. Color: trial number within block (blue=early, red=late). The RNN activity moves quickly over the stimulus decision boundary and moves slowly along the block readout direction.

State-space trajectories (Fig. 4bc) in the PCA plane (trajectories indicate how the RNN state evolves across trials in a block starting at a block change, showing a time-point per trial) show that the state jumps across the stimulus decision boundary on the timescale of trials whereas state evolves slowly and relatively steadily along the block readout, moving from the wrong block side at the start of a block (encoding the block just before the change-point) to the correct block side.

#### 3.2.2 Observations are integrated to infer stimulus side and block side

Based on state-space trajectories, we hypothesized that the RNN infers both the stimulus side and block side by integrating observations at different rates (faster for stimulus inference, slower for block inference). We confirmed this by plotting the change of RNN state along the (right-side) stimulus and (right-side) block directions, as a function of the difference in the right and left observation values 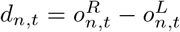 (Fig. 5ab). Both had positive slopes (0.84 for stimulus, 0.18 for block) with *p* < 1*e* − 5, confirming that evidence moves the state appropriately along the stimulus readout and block readout vectors. The respective magnitude of these two slopes (the stimulus slope is ≈ 5 times the block slope) match our expectation that stimulus side inference changes more rapidly with observations than the block side.

**Figure 5:**
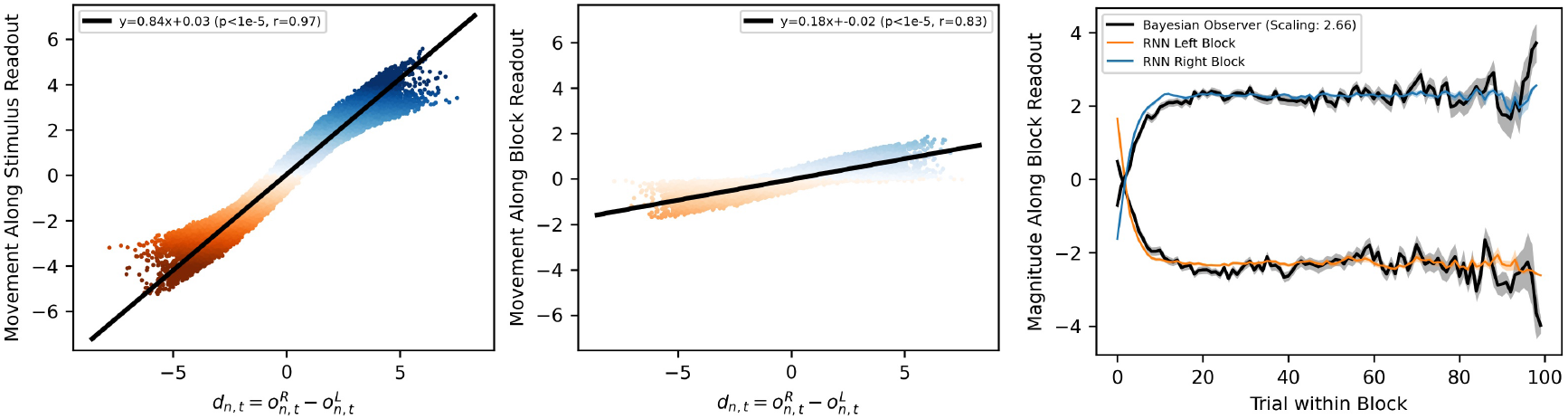
(a-b) Observations push the RNN state along the right trial readout and right block readout directions with two different amplitudes. (c) Instantaneous effects of observations are integrated along the block readout direction to encode the block side.

These instantaneous effects are integrated to infer the block side. The component of RNN activity along the right block readout vector closely matches the average block side posterior estimate of the Bayesian observer (Fig. 5c), up to an arbitrary scaling parameter that we determined through an ordinary least squares fit between the magnitude of the RNN state along the block readout vector and the Bayesian observer’s block posterior (the actor is identical to the observer in tracking the block side). This result reveals how the RNN performs efficient change-point detection of the block side.

However, when we compared the RNN’s block side belief with the Bayesian observer’s block posterior on a trial-by-trial basis, we observed a difference: The RNN block side belief, though matching the observer when averaged across trials, fluctuates more on a trial-by-trial basis (Fig. 6a). These fluctuations are driven by within-trial evidence: single right- (left-) sided trials move the RNN’s block belief to the right (left) more strongly than they move the Bayesian observer’s block posterior (Fig. 6b). This discrepancy is due to an induced dynamical coupling in the RNN between stimulus and block inference. Specifically, the RNN must update its block and stimulus beliefs simultaneously at each step and therefore cannot decouple the two inference problems, whereas both baselines decouple the two inference problems by controlling *when* information is communicated.

**Figure 6:**
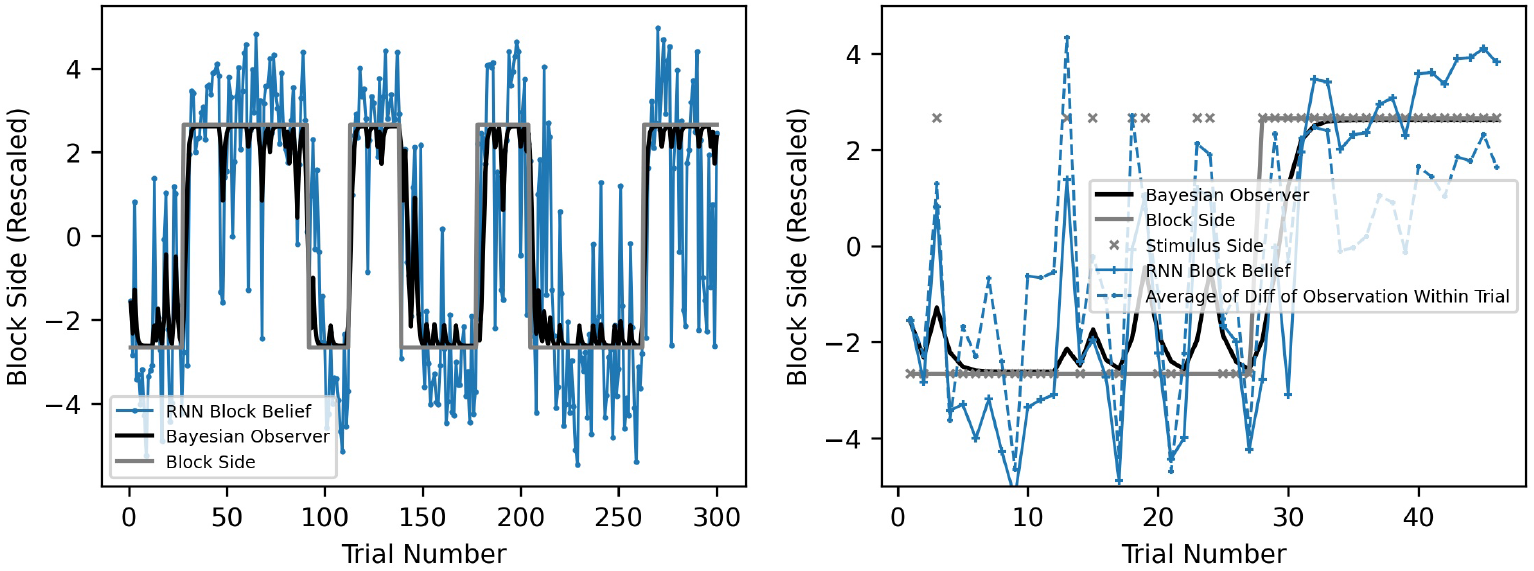
(a) RNN activity magnitude along the block readout closely matches Bayesian observer’s block posterior and the true block side. (b) Significant jumps in RNN activity magnitude along the block readout correspond to trials with large jumps in evidence, given by 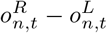.

### 3.3 RNN Mechanism

#### 3.3.1 RNN dynamics and connectivity are consistent with bistable/line-attractor dynamics

Given our hypotheses for how the dynamics of the RNN perform inference, we turned our attention to identifying the circuit mechanism(s). Ordering the hidden units using hierarchical clustering with Pearson correlation as the similarity metric revealed two clear subpopulations (Fig. 7a). Units in one subpopulation are strongly correlated with other units in the same subpopulation and strongly anticorrelated with units in the other subpopulation. Applying the same ordering to the recurrent weight matrix revealed self-excitation within each subpopulation and mutual inhibition between subpopulations (Fig. 7b).

**Figure 7:**
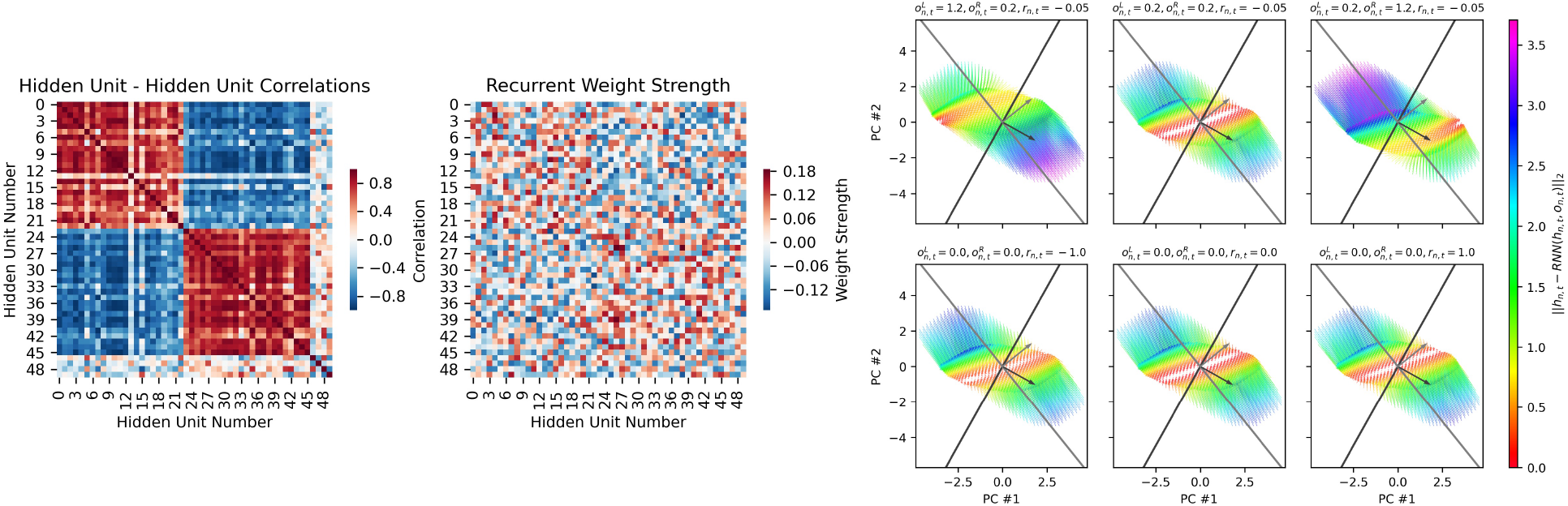
(a) Ordering RNN units based on Pearson correlation reveals two anticorrelated subpopulations. (b) Applying the same correlation-based ordering reveals self-excitatory, mutually-inhibitory connections between subpopulations. (c) RNN vector fields (evolving over one timestep) under six input conditions.

This is strongly reminiscent of circuits in the brain capable of producing 1-dimensional line-attractor dynamics or bistable attractor dynamics, depending on the strength of the excitatory and inhibitory recurrent connections [26, 18, 32, 17, 7, 23, 21]. Circuits with these dynamics have been studied in tasks involving a single variable, but not in tasks involving two (interacting) variables. We now explain how the same circuit can perform hierarchical inference on two latent variables.

Visualizing the RNN vector fields (Fig. 7c) better reveals the behavior of the system. When the stimulus is strong, the network exhibits one of two attractors, in the right-block right-stimulus quadrant or in the left-block left-stimulus quadrant. When the stimulus is absent and there is no feedback, the network exhibits a 1-dimensional line attractor. The line attractor is mainly aligned with the block readout, which allows the RNN to preserve its block side belief across trials. The persistent representation of the block side must be continuous even though the block side itself takes one of two discrete values, because the block belief is continuous-valued. The line attractor has a small projection along the stimulus readout, which translates the block belief into a stimulus prior for the next trial by biasing the RNN to select the concordant stimulus side in its decision.

Surprisingly, feedback about whether the selected action was correct has little effect (Fig. 7c). We speculate this is because the RNN’s actions are typically correct, rendering feedback less useful, and because combining feedback with the chosen action to determine the correct action requires more complex computation that is harder to learn.

#### 3.3.2 Model compression by distillation of hidden unit representations

We would like to extract a low-dimensional, interpretable model of the RNN to reveal the RNN’s effective circuit. To do so, we propose a variation of knowledge distillation [6, 4, 12] in which we train a small RNN with output states 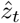 to reproduce the *hidden states* of the original RNN. This differs from conventional distillation in which the small network is trained on the *output* probabilities or logits of the original model (but a similar technique was used in BERT Transformer networks for NLP [13, 11]). We call our approach *Representation and Dynamics Distillation* (RADD). Specifically, we train the parameters *A’*, *B’* of a small RNN to recapitulate a low-dimensional projection of the original RNN’s hidden state dynamics 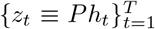, starting from initial condition 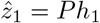, where *P* is the *M × N*-dimensional dimension-reducing projection matrix^1^. After selecting a projection P, the distilled RNN is trained on the following *L*_2_ loss using conventional methods:

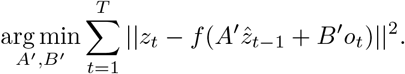

#### 3.3.3 Reduced model preserves RRN geometry and recovers meaningful parameters

The dynamics of the original RNN are well-captured by its first two principal components, suggesting that a mere 2-unit distilled RNN might suffice to capture its dynamics. Indeed, a 2-unit distilled RNN (with rows in P set to the block and trial side readout vectors; similar results are obtained with P set to the first two principal components) emulates the original RNN well: The Δ-timestep decoherence in state is the same across three systems, ||*h*_*t*+Δ_ − *h_t_*||, ||*z*_*t*+Δ_ − *z_t_*|| (Fig. 8a). States in the distilled RNN evolve in a qualitatively similar way across the trial and block boundaries over multiple trials as the original RNN (Fig. 8b). Moreover, depending on the magnitude of the distilled system’s readout vector (a free parameter), the distilled system can slightly outperform the full RNN (distilled 86.87%, full 85.50%) on the task. The distilled 2-unit RNN recognizes blocks in the same way as the original RNN, whereas a 2-unit RNN trained directly on the task itself fails to recognize blocks (Fig. 8c) despite being trained for four times as many gradient steps.

**Figure 8:**
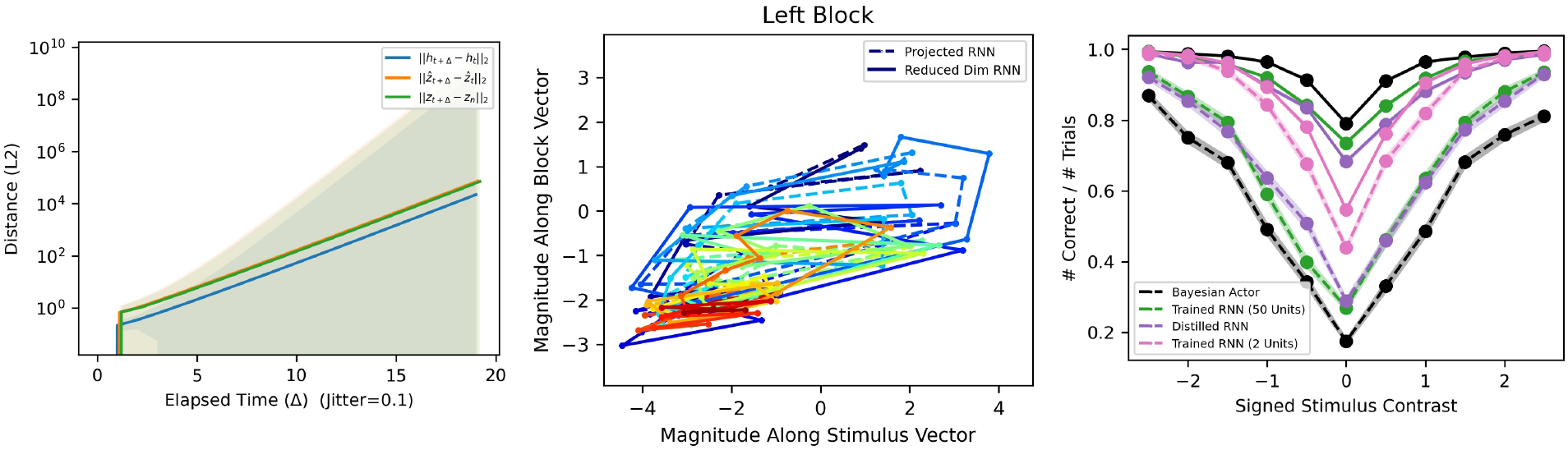
(a) Distance decoheres at the same rate in the three state spaces: RNN, projected RNN, distilled RNN. Projected RNN and distilled RNN have nearly identical values (horizontal displacement added to make both visible). (b) Distilled RNN state space trajectories closely match projected RNN trajectories. (c) Comparative performance of 50-unit original RNN, 2-unit distilled RNN, and 2-unit task-trained RNN.

The distilled RNN, whose units correspond to stimulus and block side beliefs, has sensible parameters:

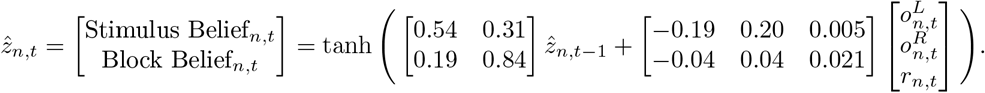

The recurrent weights show that the stimulus belief and block belief reinforce one another, and both decay to 0 without input observations. The input weights show that observations drive the stimulus and block side beliefs in a common direction, but that the movement caused by a single observation is 5 times greater along the stimulus direction than the block direction. The state space trajectories (Fig. 8b) visually agree with this intuition: each left-to-right (stimulus side) movement corresponds to a small up-right (block side) movement. Further, the feedback input receives negligible weighting, consistent with our earlier observation.

## 4 Discussion

In conclusion, RNNs attain near-optimal performance on a hierarchical inference task, as measured against Bayesian observers and actors that have full knowledge of the task. We have characterized the representations, dynamics, and mechanisms underlying inference in the RNN. In future work, we will leverage these models, together with work being developed by others, to better understand mouse behavior and neural representations. We expect it will be fruitful to explore RADD in the context of reinforcement learning [24, 9].

## Footnotes

1 We assume that the *N*-dimensional states of the original RNN lie in a *D*-dimensional linear subspace ℝ^*D*^ ⊂ ℝ^*N*^. The projection *P* can be selected such that *D* ≤ *M* ≤ *N* and such that its nullspace does not intersect this *D*-dimensional subspace. A good choice for *P* are the top principal vectors of the original RNN states.

